# GraphAligner: Rapid and Versatile Sequence-to-Graph Alignment

**DOI:** 10.1101/810812

**Authors:** Mikko Rautiainen, Tobias Marschall

## Abstract

*Genome graphs* can represent genetic variation and sequence uncertainty. Aligning sequences to genome graphs is key to many applications, including error correction, genome assembly, and genotyping of variants in a pan-genome graph. Yet, so far this step is often prohibitively slow. We present GraphAligner, a tool for aligning long reads to genome graphs. Compared to state-of-the-art tools, GraphAligner is 12x faster and uses 5x less memory, making it as efficient as aligning reads to linear reference genomes. When employing GraphAligner for error correction, we find it to be almost 3x more accurate and over 15x faster than extant tools.

**Availability Package manager:** https://anaconda.org/bioconda/graphaligner and source code: https://github.com/maickrau/GraphAligner

## 1 Background

Graphs provide a natural way of expressing variation or uncertainty in a genome [1, 2]. They have been used for diverse applications such as genome assembly [3, 4, 5], error correction [6, 7, 8], short tandem repeat genotyping [9], structural variation genotyping [10] and reference-free haplotype reconstruction [11]. With the growing usage of graphs, methods for handling graphs efficiently are becoming a crucial requirement for many applications.

Sequence alignment is one of the most fundamental operations in bioinformatics and necessary for a wide range of analyses. Aligning a sequence to a sequence is a well studied problem with many highly optimized tools [12, 13, 14, 15]. In contrast, aligning sequences to graphs is a newer field and practical tools only start to emerge, where most of the existing tools are specialized for one purpose such as error correction [6, 7, 8], or hybrid genome assembly [4]. The VG toolkit [16] provides a set of general-purpose tools to work with genome graphs. Although VG is capable of mapping long reads to graphs, it was tuned for aligning short reads, leading to slow runtimes for long read alignment. In summary, there is presently a lack of general-purpose tools for aligning long third-generation sequencing reads to graphs. Given the wide range of applications, including sequence assembly, error correction, and variant calling, and the steep decline in prices for long read sequencing, closing this gap is critical.

Outside of the bioinformatics community, an algorithm for aligning sequences to an arbitrary graph with unit costs was already discovered in 2000 in the context of hypertext searching by Navarro [17]. An important property of Navarro’s algorithm is that the runtime depends only on the number of nodes and edges and the length of the query sequence. Thus complex cyclic graphs are (asymptotically) just as easy as simple linear graphs of the same size. Recently it was proven that the runtime of Navarro’s algorithm is in fact optimal unless the strong exponential time hypothesis is false [18]. In 2002, *partial order alignment* [19] (POA), a special case of Navarro’s algorithm for acyclic graphs, was published for multiple sequence alignment. Although POA is defined only for acyclic graphs, it can be extended to cyclic graphs by unfolding cyclic components, which is the approach taken by the VG toolkit [16] and ExpansionHunter [9]. The practical efficiency of this unfolding depends on the read length and the graph topology and complex cyclic areas can lead to very large unfolded graphs [20]. V-Align [20] aligns to cyclic graphs but its runtime depends on the graph’s feedback vertex set size. Some tools use heuristic approaches for aligning to de Bruijn graphs using depth-first search [6, 21, 8]. Navarro’s algorithm has recently been generalized to arbitrary costs as well [22]. Our previous work [23] combined Navarro’s graph alignment algorithm with Myers’ bit-parallel algorithm [24], leading to speedups in practice between 5x-20x, but this algorithm is designed to compute the full dynamic programming table, making it unsuitable for aligning many reads to a large reference graph.

In contrast, most practical tools use a *seed-and-extend* strategy. Seeding depends on finding matches between the read and the graph, and necessitates indexing the graph in some manner. Although asymptotically optimal algorithms for graph alignment are known, the lower bound for indexing a graph is currently unknown. K-mer based indices have been used in many de Bruijn graph alignment tools [6, 21, 25]. The Positional Burrows-Wheeler transform [26] is a method for indexing multiple sequence alignments between genomes, which can be viewed as a special class of graph genomes. Indexing variation graphs is challenging because the number of possible paths can be exponential in the number of variants encoded. Typical approaches to handle this problem are to index only some of the variation by limiting the indexed paths either heuristically [16, 27, 28] or by using panels of known haplotypes [29, 30]. A recent method avoids the exponential blowup by dynamically indexing the graph and the reads, thereby exploiting that there can be exponentially many paths in the graphs, but not in the set of reads to be queried [31].

*Contributions*. Here, we provide the first algorithm for *banded* sequence-to-graph alignment that scales to align noisy long reads to de Bruijn graphs of whole human genomes. We also apply a simple minimizer [32] based seeding method which exploits the fact that long reads almost always span simple areas of the genome, unlike short reads which are more prone to being entirely embedded within a variation-rich area.

We describe our sequence-to-graph long read alignment tool GraphAligner. GraphAligner is designed to work with arbitrary graphs instead of specializing for one type of graph. We compare GraphAligner to two well-optimized tools for linear alignment [13, 14], and to the vg toolkit [16] for aligning to variation graphs. To show how better alignment methods improve downstream applications, we present a pipeline for error correcting long reads based on graph alignment, which we compare to existing methods based on the same principle. Although using a similar process as existing tools, the better alignment strategy leads to an order of magnitude speedup and error rates around one third of the current state-of-the-art for whole human genome data.

## 2 Results

### 2.1 Comparison to linear aligners

Regular sequence-to-sequence alignment is a special case of sequence-to-graph alignment, where the graph consists of a linear chain of nodes. We compare GraphAligner to two well-optimized sequence-to-sequence aligners, minimap2 [13] and BWA [14]. We use a PacBio dataset^[1]^ from human individual HG00733, randomly subsampled to 15x coverage. For minimap2 and BWA, we give the GRCh38 reference genome as-is. For GraphAligner, we first split the reference at each non-ATCG character, and then create a graph which has each contig of the reference as a node, without any edges. Aligning to this graph is equivalent to aligning to the linear reference. To measure the alignment quality, we measure the amount of sequence aligned and the error rate of the alignments.

Table 1 shows the results. There are small differences in the amount aligned and the error rate, showing that the aligners have slightly different tradeoffs between sensitivity and specificity, but all three aligners roughly agree on the amount aligned and the error rate. On runtime, there is a large difference between BWA and the two other aligners, about 90x between minimap2 and BWA, and 85x between GraphAligner and BWA. GraphAligner is slightly slower than minimap. Overall the results show that GraphAligner is competitive with the state of the art for linear alignment.

**Table 1.** Results of the linear comparison experiment. PacBio reads were aligned to the GRCh38 reference genome with minimap2, BWA mem and GraphAligner.

| Aligner | Aligned (bp, | %) | Error rate (%) | CPU-time<br>(HH:mm:ss) | Peak memory (Gb) |
| --- | --- | --- | --- | --- | --- |
| Original | 48 825 600 292 | 100 | - | - | - |
| minimap2 | 47 669 242 462 | 97.6 | 14.1986 | 19:00:22 | 43.7 |
| BWA mem | 47 174 318 479 | 96.6 | 13.8979 | 1752:17:25 | 13.1 |
| GraphAligner | 47 129 226 576 | 96.5 | 13.6979 | 20:36:10 | 29.8 |

### 2.2 Variation graph

In this experiment we built a variation graph of the human chromosome 22 and compared GraphAligner and vg [16] on it. To build the graph, we took the GRCh38 reference and variants [33] called by the Human Genome Structural Variation Consortium [33], and used vg to build the graph from the reference and the variants. We first randomly subsampled the reads^[2]^ to 15x coverage. Then we selected the reads by first aligning the subsampled reads to the GRCh38 reference with minimap2, then selecting reads which have an alignment to chromosome 22 containing at least 70% of the read, and no alignments to other chromosomes.

Table 2 shows the results. vg reported all base pairs as aligned while GraphAligner clipped the read ends when the alignment score became too poor, which resulted in 98.9% of all base pairs being aligned. GraphAligner’s runtime and peak memory includes both indexing and alignment. Despite including the indexing phase, we see that GraphAligner is almost nine times faster than vg’s mapping phase. When including vg’s indexing as well, GraphAligner is over twelve times faster than vg. Peak memory use is five times smaller.

**Table 2.** Results of the variation graph experiment. PacBio reads were aligned to a chromosome 22 variation graph using both GraphAligner and vg.

| Aligner | Aligned (bp, | % | CPU-time<br>(HH:mm:ss) | Peak memory (Gb) |
| --- | --- | --- | --- | --- |
| Original | 413 001 450 | 100 | - | - |
| vg index | - | - | 0:36:43 | 9.3 |
| vg map | 413 001 450 | 100 | 1:34:40 | 3.4 |
| GraphAligner | 408 662 977 | 98.9 | 0:10:32 | 1.7 |

### 2.3 Error correction

We have implemented a hybrid error correction pipeline based on sequence-to-graph alignment. Aligning reads to a de Bruijn graph (DBG) is a method of error correcting long reads from short reads [6, 7]. The idea is to build a DBG from the short reads and then find the best alignment between the long read and a path in the DBG. The sequence of the path can then be used as the corrected long read.

Zhang et al. [34] performed an evaluation of 16 different error correction methods. Based on their results, we chose FMLRC [8] as a fast and accurate hybrid error corrector for comparison. We also compare to LoRDEC [6] since our pipeline uses the same overall idea as they do.

LoRDEC [6] builds a de Bruijn graph from the short reads, then aligns the long reads to it using a depth-first search and uses the path sequence as the corrected read. FMLRC [8] also aligns the reads to a graph, except instead of building one de Bruijn graph, it uses an FM-index which can represent all de Bruijn graphs and dynamically vary the k-mer size. FMLRC then corrects the reads in two passes, using different k-mer sizes. Our error correction pipeline is similar to LoRDEC. Figure 1 shows the pipeline. We first self-correct the Illumina reads using Lighter [35], then build the de Bruijn graph using BCalm2 [36], align the long reads using GraphAligner with default parameters and finally extract the path as the corrected read.

**Figure 1.**
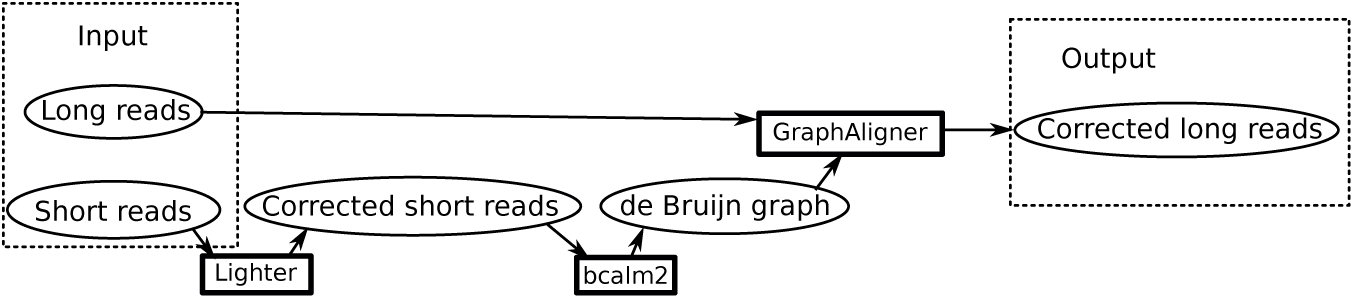
Overview of the error correction pipeline. The circles represent data and the rectangles programs.

Due to fluctuations and biases of Illumina coverage, some genomic areas are impossible to correct with short reads even in principle. Our pipeline has two modes: either we output the full reads, keeping uncorrected areas as is; or clipped reads, which remove the uncorrected areas and split the read into multiple corrected subreads, if needed. In the results, we present the full reads as “GraphAligner”, and the clipped reads as “GraphAligner-clip”. We similarly report “LoRDEC” as full reads and “LoRDEC-clip” as clipped reads. FMLRC does not offer an option to clip the reads so we report only the full reads.

To evaluate the results, we use the evaluation methodology from Zhang et al. [34]. The long reads are first corrected, and then the evaluation pipeline is run for both the raw reads and the corrected reads. The first step of the evaluation is removing reads shorter than 500 bp. Note that the reads are removed during the evaluation step, that is, they are corrected in the initial correction step and different reads may be removed in the uncorrected and corrected sets. After this, the remaining reads are aligned to the reference genome. The alignment yields several quality metrics, including number of aligned reads and base pairs, read N50, error rate and genomic coverage. Here, we report error rate as given by samtools stats instead of alignment identity. Resource consumption is measured from CPU time and peak memory use. We use the *E. coli* Illumina+PacBio dataset (E. coli, called D1-P + D1-I by Zhang et al.) and the *D. melanogaster* Illumina+ONT dataset (Fruit fly, called D3-O + D3-I by Zhang et al.) from Zhang et al. [34]. In addition, we use whole human genome PacBio Sequel^[3]^ and Illumina^[4]^ data from HG00733, randomly subsampled to 15x coverage for PacBio and 30x for Illumina. We use the diploid assembly from [33] as the ground truth to evaluate against for HG00733. We did not include LoRDEC in the fruit fly or HG00733 experiments as the results in [34] show that FMLRC outperforms it in both speed and accuracy. Although we use the same evaluation method, our results are slightly different. This is due to two factors: First, Zhang et al. use LoRDEC version 0.8 with the default parameters, while we use version 0.9 with the parameters suggested for *E. coli* in the LoRDEC paper [6]. Second, Zhang et al. use FMLRC version 0.1.2 and construct the BWT with msBWT [37], while we use version 1.0.0 and construct the BWT with RopeBWT2 [38] as recommended by the FMLRC documentation.

Table 3 shows the results. The amount of aligned sequence is similar in all cases. The amount of corrected sequence is lower than the original in PacBio datasets and higher in the ONT dataset. This is consistent with the observation that insertion errors are more common than deletions in PacBio and vice versa for ONT [39]. The number of reads is noticably higher and the N50 is lower for the clipped modes for both LoRDEC and GraphAligner, showing that most reads contain uncorrected areas and clipping the reads reduces read contiguity. In addition, the fruit fly and human experiments show that clipping the reads significantly reduces the genome fraction covered by the reads. The clipping is more pronounced in the more complex genomes, with the reads in the whole human genome dataset being on average cut into four pieces, around 5% of the genome lost due to clipping and a large reduction in read N50. We see that GraphAligner is almost 23x faster and 2.7x more accurate than LoRDEC for *E. coli*. GraphAligner is over eight times faster than FMLRC in all datasets. When not clipping reads, GraphAligner’s error rate is slightly worse then FMLRC for *E. coli* (0.52% vs. 0.30%), but substantially better for *D. melanogaster* (1.4% vs. 2.3%) and human (2.6% vs. 7.1%). For the human genome HG00733, GraphAligner hence produces almost three times better error rates while the runtime is over 15x times faster.

**Table 3.**
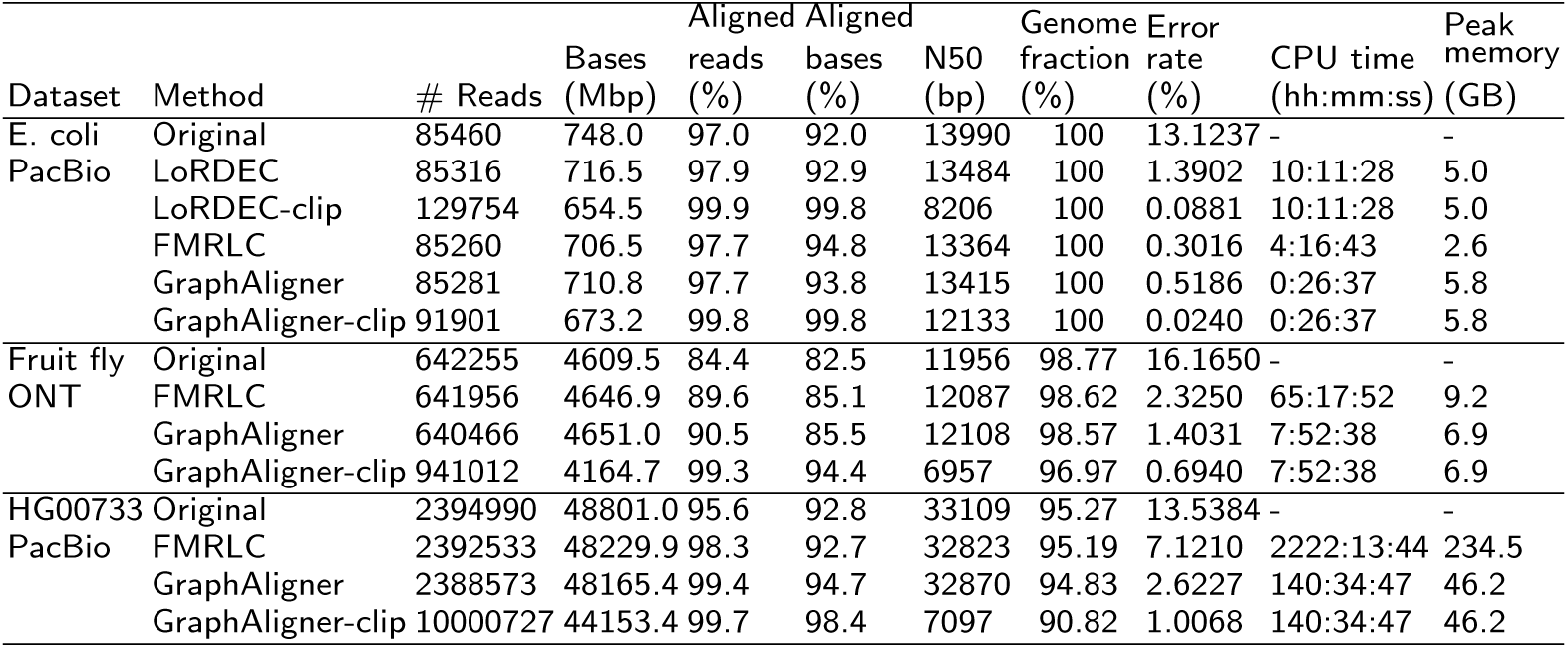
Results of the error correction experiment. Reads shorter than 500 base pairs are discarded. The remaining reads were aligned to the reference using minimap2 [13] and the statistics were given by samtools [40] stats, except N50 which is calculated by a script from Zhang et al [34] and resource use which are measured by “/usr/bin/time -v”.

Our pipeline is a large improvement in runtime over the state-of-the-art. The error rates are competitive for simpler genomes and significantly better for more complex genomes. We hypothesize that the two-pass method used by FMLRC can in principle enable better correction than a single k-mer size graph, but FMLRC’s performance with the larger genomes is limited by their alignment method, while GraphAligner can handle the more complex genomes. When using the clipped mode, that is, when only considering parts of the reads that have been corrected, the accuracy in the corrected areas can approach or exceed the accuracy of short reads. This emphasizes the value of this clipped mode to users. The main source of errors are in fact uncorrected areas without sufficient short read coverage.

## 3 Discussion

We have presented GraphAligner, a tool for aligning long reads to sequence graphs. Although GraphAligner is designed for graphs, it can also align to trivial linear graphs, and the performance is comparable to state of the art linear mappers. However, for linear alignment we recommend using linear mappers due to the different data formats used for graph alignment (GAM[16] instead of BAM[40]), which standard downstream tools normally do not accept. In non-trivial variation graphs, GraphAligner outperforms vg by a factor of 12 in runtime.

GraphAligner is presently geared towards aligning long reads, which was our focus due to the absence of methods for this. The current seeding strategy can systematically fail to handle short reads in variation-dense regions. However, the core algorithmic components of GraphAligner could likely be used to also align short reads. To this end, we plan to integrate GraphAligner with PSI [31], a novel seeding approach that we developed recently to facilitate efficient and full-sensitivity seed finding across node boundaries.

As sequence alignment is a very fundamental operation and long reads are rapidly becoming more affordable to produce, we anticipate that GraphAligner will be used widely and will improve the performance and runtime of many downstream applications. Here, we have shown one example of this with our error correction experiment, where our pipeline improves on the state of the art and enables correcting long reads in mammalian scale genomes to high accuracy. It would be possible to combine GraphAligner’s alignment with the FM-index based graph as used by FMLRC, which might yield an error correction pipeline as fast as and more accurate than our current results, which is an interesting avenue for future developments. Other applications such as graph-based hybrid genome assembly also align reads to a graph, either explicitly [4] or by reducing the problem to sequence-to-sequence alignment [5]. It is likely that improved alignment methods will lead to improved results here as well, and we are currently investigating this further. Lastly, GraphAligner might enable scaling the haplotype-resolved genome assembly method that we demonstrated for yeast genomes [11] to mammalian genomes.

## 4 Conclusions

We have implemented the sequence-to-graph alignment tool GraphAligner. As genome graphs become more common, efficient methods for aligning reads to genome graphs become more important. GraphAligner is competitive with well-optimized linear aligners when aligning to a linear genome, and outperforms existing graph alignment tools 12x in runtime. We have implemented a long read error correction pipeline using GraphAligner, and show that the method outperforms the current state-of-the-art, with an almost 3x improvement in error rate and over 15x improvement in runtime for whole human genomes.

## 5 Methods

Figure 2 shows an overview of GraphAligner. One IO thread reads sequences, which are passed to an arbitrary number of worker threads. Each worker thread aligns reads one at a time. The alignment algorithm uses a seed-and-extend method. Seeds are found by matching the read with the node sequences, and then extended independently of each others with a bit-parallel banded dynamic programming algorithm. Finally the primary and supplementary alignments are selected and passed to a second IO thread, which writes the results to a file.

**Figure 2.**
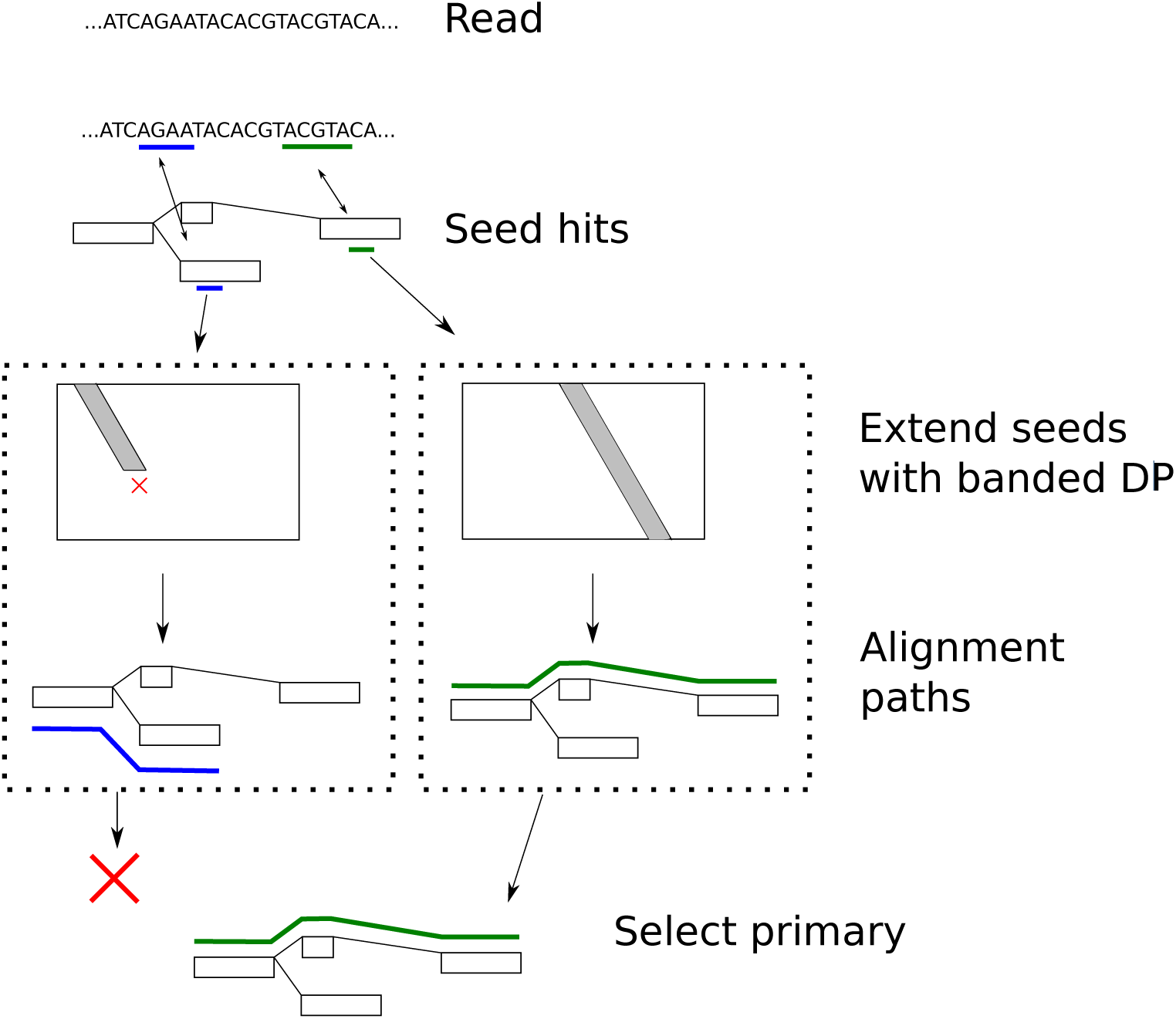
Overview of GraphAligner. Reads are aligned independently of the other reads. Seed hits are found by matching the sequence of the read to sequences inside nodes (small blue and green bars). Seed hits are then extended independently of each other (small dotted boxes) with a banded dynamic programming algorithm, using Viterbi’s algorithm to decide when to clip the alignment (red X). Each seed hit can result in an alignment (blue and green paths). Alignments that overlap with an another, larger alignment are classified as secondary. Secondary alignments are discarded by default (red X) but can be included in the output with an optional parameter. The output is then written to a file either as alignments or corrected reads.

### 5.1 Data formats

We designed GraphAligner to use the most common file formats, and specifically be interoperable with vg [16] to leverage existing graph-based operations and pipelines. Graphs are inputed either in the binary vg graph format [16] or the human-readable graphical fragment assembly (gfa) format [41]. By allowing gfa, GraphAligner is moreover able to handle graphs with overlapping node labels, which is presently not supported by the vg file format. Reads are inputed in fasta or fastq, and optionally gzip-compressed. Alignments are outputed in vg’s binary *gam* format, a generalization of SAM/BAM format [40] to graphs. Alignments can also be out-puted in an equivalent human-readable JSON format.

### 5.2 Graph model

GraphAligner inputs *bidirected graphs* [16], which are capable of representing genome graphs commonly used in bioinformatics, including de Bruijn graphs [42, 36], assembly graphs [43, 3, 44], pan-genomes [1], and variation graphs [2, 16]. Bidirected graphs model the double-stranded nature of DNA. The sequence is stored in the nodes, which can be traversed in two directions; either left to right with the node label, or right to left with the reverse complement of the label. The edges connect to either the *left end* or the *right end* of a node. A path through a bidirected graph enters a node from one end, traverses through the node, and then leaves via an edge in the opposite end.

The bidirected graph is first converted into a directed node-labeled graph which we call the *alignment graph*, with a mapping between the bidirected graph and the alignment graph. The read is then aligned to the alignment graph, and the mapping is used to convert the alignment back into the bidirected graph.

The bidirected graph allows an overlap between edges, representing for example overlapping *k* − 1-mers of a de Bruijn graph, or the read overlap in an assembly graph. Here, we consider the edges to be labeled by the number of overlapping nucleotides. When traversing via an edge with an overlap of *n* nucleotides, the path must skip the first *n* nucleotides of the target node. The overlaps can also vary between edges. Edge overlaps are handled by chopping the node into pieces at each overlap boundary. The alignment graph then has edges connecting the end of a node to the chopped boundary of the neighbor. This allows a path that ends at one node to enter the neighboring node without traversing the overlap twice. Figure 3 shows an example of the edge chopping for edges with variable overlaps. In addition to this, nodes longer than 64 base pairs will be chopped to multiple nodes, each containing up to 64 base pairs.

**Figure 3.**
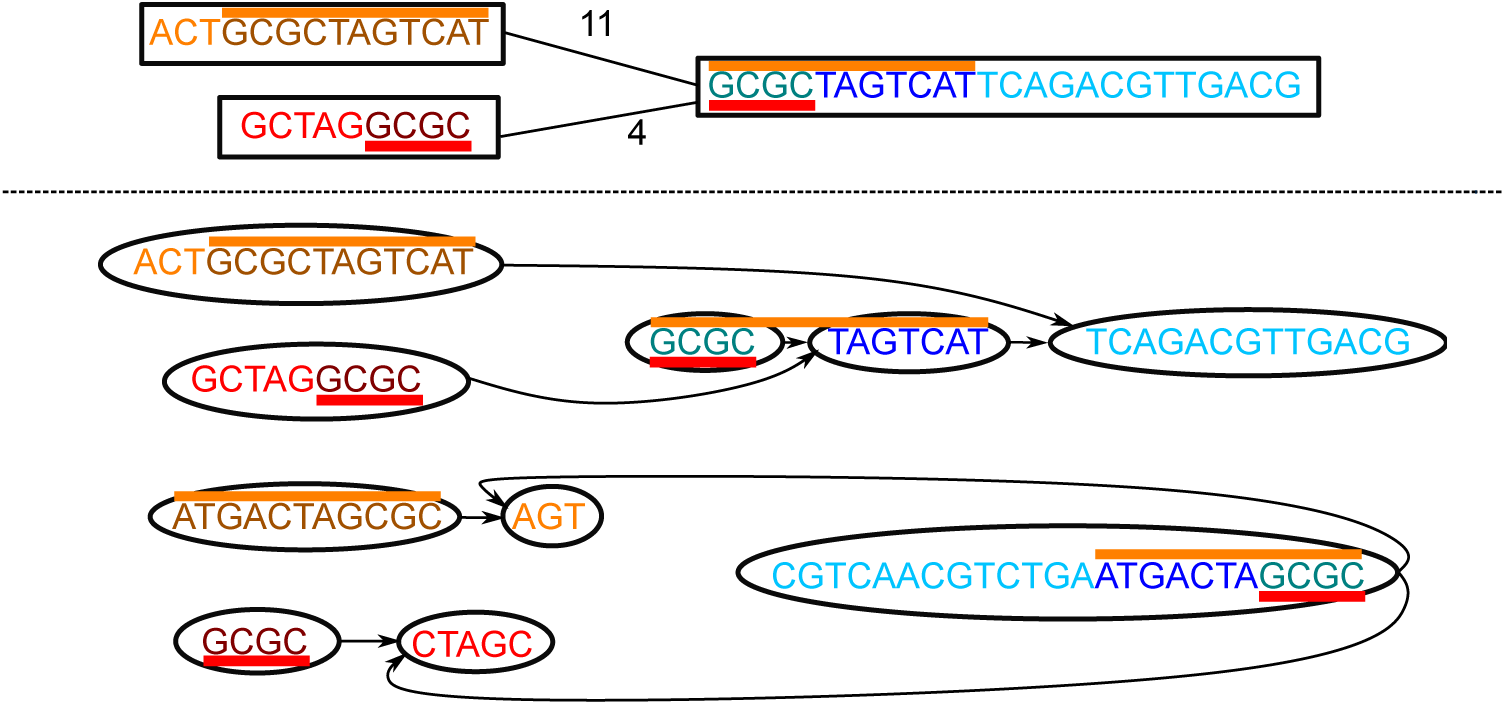
Converting a bidirected graph with variable edge overlaps to an alignment graph. Top: a bidirected graph with three nodes. The edges are labeled by their overlap. The red colored bars represent the same sequence, which should not be duplicated during traversal. Similarly, the orange colored bars represent the same sequence. Bottom: the alignment graph created from the top graph. The colors of the base pairs show how they match between the two graphs, with each sequence in the original graph represented by the same color in the alignment graph twice, once for the forward strand and once for the reverse complement. Similarly to the bidirected graph, the red and orange bars represent the same sequences. There are two subgraphs, one representing the forward traversal (top) and one the backward traversal (bottom) with reverse complemented node labels. Each edge introduces a breakpoint in the target node, splitting the node at the boundary of the overlap. The alignment graph then connects the ends of the overlap such that the overlapping sequence is only traversed once.

A node in the bidirected graph with *l* nucleotides adds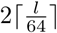he alignment graph,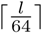ward traversal and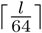kward traversal, and each edge can split up to two nodes and add up to four edges in the alignment graph. The number of nucleotides in the alignment graph is exactly twice the number of nucleotides in the bidirected graph. Therefore the transformation produces an alignment graph whose size is within a constant factor of the bidirected graph.

Both the read and the graph are allowed to contain ambiguous nucleotides (B, R, N, etc.) The alignment extension considers two ambiguous nucleotides a match if any of the possible nucleotides match; eg, R (A or G) matches W (A or T) because both of them could be A, but R (A or G) does not match Y (C or T) because there is no overlap. Only the non-ambiguous characters A, T, C and G are used for seeding.

### 5.3 Seeding

The first part of the seed-and-extend algorithm is finding seed hits. Here, we define seeds as exact matches between a read and a node sequence, but other definitions exist in the literature.

Typical alignment approaches [13] *chain* seeds to find the approximate position of the alignments. For linear sequences, seed chaining is solved with the *co-linear chaining* problem that exploits the fact that calculating the distance between seeds in a linear sequence is trivial. However, for graphs, the distance between seeds can be ambiguous as there are multiple paths connecting the seeds, and finding the distance in a graph is computationally more expensive than in a linear sequence [45]. For these reasons, GraphAligner uses a different approach: Seeds are not chained, but instead each seed is extended separately, starting with the most promising seed. The alignment algorithm used for this extension step decides which paths to explore and when to end the alignment (detailed in Section 5.4). Seeds included in alignments from previously explored seeds are skipped. This strategy leads to relaxed requirements for the seeding algorithm; instead of requiring seeds along the entire valid alignment, it is enough to find at least one valid seed per read from which to extend.

GraphAligner uses a simple method for transforming text matching in graphs to text matching in strings. Instead of matching reads to paths in the graph, reads are matched to node sequences in the graph. The nodes can be treated as a collection of strings which enables using efficient string matching algorithms. Figure 4 shows an example of matching a read to nodes in a graph. Note that we use the node sequences from the original bidirected graph, not from the directed alignment graph. Reverse complement matches are also allowed.

**Figure 4.**
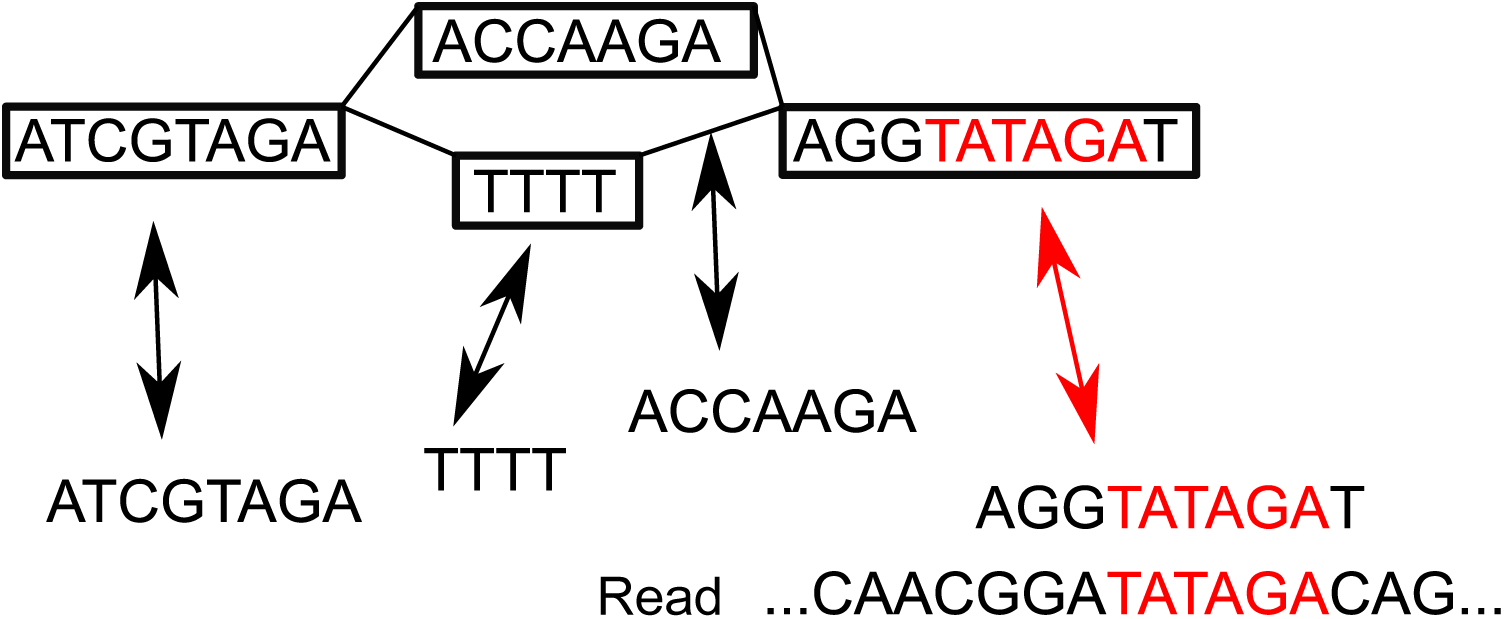
Seeding. Top: A graph with four nodes. Middle: The node sequences are extracted from the nodes. The arrows represent a mapping between the strings and nodes. Bottom: A read. Highlighted in red: Matches between the read and a string are transformed into a match inside a node using the mapping.

This approach finds only seed hits which are entirely contained in a node. For the special case of de Bruijn graphs, we hence finds hits of length up to *k* due to the overlap between the nodes. However, in general it misses seeds which cross a node boundary. We have noticed that in practice this is not an important limitation for long reads, since the read almost always crosses linear parts of the graph which can be used for finding at least one seed hit. For example, in the chromosome 22 variation graph experiment, 99.8% of reads had at least one seed hit and 99.99% of the base pairs were contained in a read with at least one seed hit.

The default method for finding matches is by using *minimizers* [32]. A window of *w* base pairs is slid through the text and the smallest *k*-mers of each window according to a hash function are picked as the minimizers. For the seeding string, the minimizers are kept in a hash table that keeps the count and locations for each *k*-mer. For the read, the minimizers are further split into *c*-base pair *chunks*, and the *n* least frequently appearing *k*-mers in each chunk are kept as seed hits. The chunking ensures that there are seed hits along the entire read, and the least frequently appearing *k*-mers are more likely to be correct seeds. Figure 5 shows this process. The default values use *k* = 19, *w* = 30, *n* = 5 and *c* = 100. These values were chosen empirically after aligning reads to a whole human genome variation graph with different parameter sets. The index is implemented with succinct data structures from the SDSL library [46] and minimal perfect hashing from BBHash [47].

**Figure 5.**
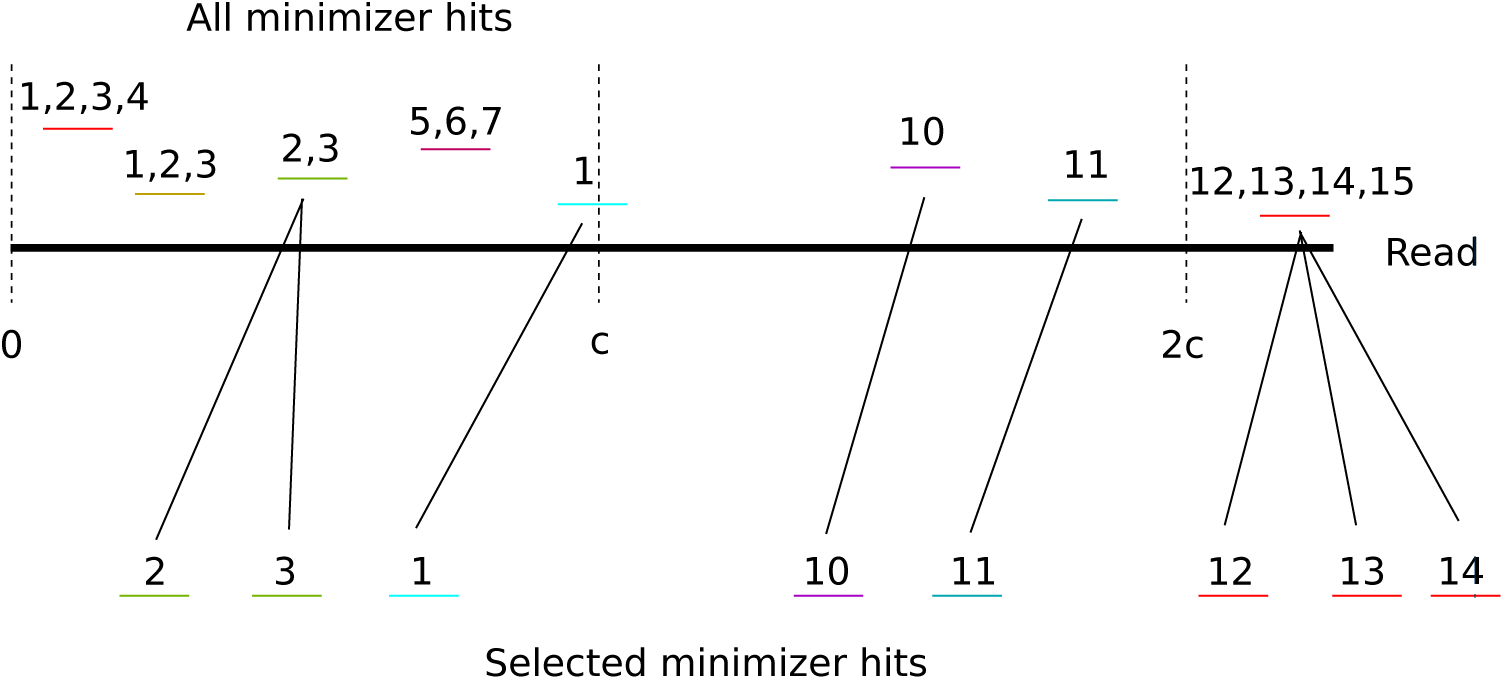
Minimizer seed selection. The read is represented by the thick black line and the minimizer hits are represented by the small colored lines. The numbers above a minimizer represent positions in the graph that match the minimizer. The frequency of a minimizer is the count of positions in the graph that match the minimizer. The read is divided into chunks of up to *c* base pairs (dotted lines), and minimizers are assigned into the chunks according to their position in the read. In each chunk, *n* (here *n* = 3) minimizer matches are selected from the least frequent minimizers, with ties broken arbitrarily. The thin black lines show how the selected minimizers correspond to the minimizer hits.

In addition to the built-in seeding methods, seeds can be inputed from a file, allowing an arbitrary external method to be used for seeding. The seeds must then be provided in GAM format [16], containing a position in the read, a position in the graph and a match length. The match length is only used for deciding an ordering for the seeds, and does not need to actually correspond to the length of the match.

A heuristic method is used to decide which seeds are extended. Starting from the least frequent seed hit, the alignment is extended as far as possible. Then, for each more frequent seed, if the seed is inside a part of the read which has already been aligned, the seed is discarded. There is an optional switch to extend those seeds as well, which is off by default.

Finally, GraphAligner has a mode for aligning without seeds. In this case, the extension algorithm is initialized with the entire first row of the dynamic programming table being considered and then proceeding as usual (see next section for details). In this way, the alignment algorithm would implicitly scan the whole graph. The runtime is dependent on the graph size, so this mode is only practical for graphs up to a few million base pairs in size.

### 5.4 Extension

GraphAligner uses a dynamic programming (DP) algorithm to extend the seeds. The starting point of the DP is the well known Needleman-Wunsch algorithm for sequence alignment [48]. This algorithm has been generalized to sequence-to-graph alignment by Navarro [17]. In a previous work [23] we further generalized Myers’ bit-parallel method [24] to sequence-to-graph alignment to improve the runtime.

In short, the algorithm calculates a *DP matrix* whose scores describe the edit distance of an alignment ending at a specific position in the read and a specific position in the graph. The calculation proceeds in a sliced manner, first calculating a horizontal slice of the topmost 64 rows, then calculating the next topmost slice and so on. For details on how to calculate the DP matrix for graphs in a bit-parallel manner, we refer the reader to [23]. In the following focus on describing the extensions over this previous work that were necessary to make GraphAligner scale to large graphs: first, a faster algorithm for merging bitvectors; second, how to apply *banded alignment* [49] to graphs, reducing the area in the DP matrix which needs to be calculated and greatly reducing runtime and memory use; and third, how to efficiently store a partial DP matrix of a graph.

### 5.5 Bit-parallel operations

The DP extension algorithm requires merging bitvectors at nodes with an in-degree of at least two. In our previous work [23] we described an *O*(log *w*) algorithm for merging two bitvectors. We have refined this operation further and created an algorithm which is much faster in practice but with a theoretically slower runtime of *O*(*w*). In practice, the *O*(*w*) algorithm takes on average around 50 instructions per merge, while the *O*(log *w*) algorithm takes on average around 300 instructions per merge for 64-bit bitvectors. The code and detailed explanation of the merging algorithm is in the supplementary material.

### 5.6 Banded alignment on graphs

In sequence-to-sequence alignment, banded alignment [49, 50] is a technique for speeding up the alignment while guaranteeing that the optimal alignment is still found as long as the number of errors is small. The idea is that given a start position of the alignment and a maximum edit distance, a diagonal parallelogram is selected, and the DP matrix is calculated only inside the parallelogram [50]. Given a banding parameter *b*, the width of the parallelogram is 2*b* and the optimal alignment is guaranteed to be found if it has at most *b* errors. The runtime of the alignment no longer depends on the size of the reference, leading to a large speedup.

The parallelogram technique cannot be used in graphs due to the non-linear structure. At each fork, the parallelogram should continue to both paths. This would mean that the size of the band could grow very large and the bookkeeping involved in tracking the band would introduce heavy overhead, possibly exponential to the size of the graph.

Instead we introduce a novel dynamic banding approach based on the scores in the DP matrix. The principle is that for each row, we define a cell to be inside the band if its score is within *b* of the minimum score for that row. This handles arbitrary graph topologies without any extra bookkeeping or special cases. Figure 6 shows an example. Since the band depends on the minimum score in a row, we do not initially know which parts of the DP matrix are included in the band. Instead, we “discover” the minimum score and the edges of the band as we calculate the DP matrix.

**Figure 6.**
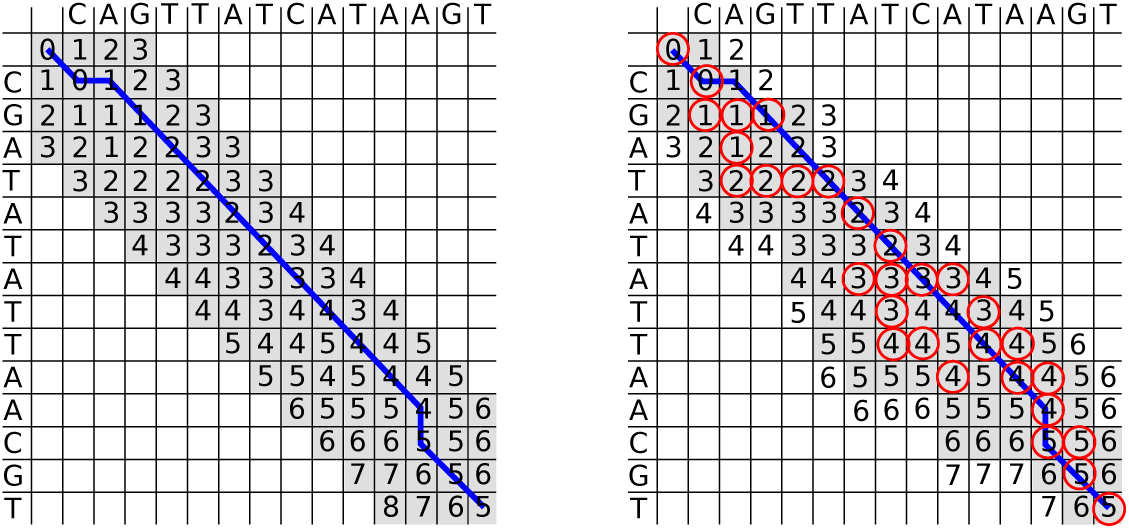
Left: regular banded alignment with *b* = 3. The reference is on top and the query on the left. The gray cells are inside the band and are calculated. The blue line shows the traceback of the optimal alignment. Right: score based banding with *b* = 1. The reference is on top and the query on the left. The gray cells are inside the band and the blue line is the traceback. The red circled cells are the minimum for each row, which are discovered during the calculation of the matrix and define whether a cell is inside the band or not; a cell is inside the band if its score is within *b* of the minimum score in the same row. The cells with a number on a white background are calculated to discover the end of the band, but they are not inside the band and are ignored when calculating the next row. The band can wander around the DP matrix and change size, automatically spreading wider in high error areas and narrower in low error areas. Note that the score based banding parameter is 1 in comparison to 3 in the regular banding to the left. The implementation uses a coarser band of 64 x 64 blocks instead of individual cells.

Figure 7 shows how the dynamic score-based banding handles different topological features. At each fork, the band spreads to all out-neighbors. This explores the different paths the alignment could take, while the score comparison implicitly limits how far the exploration proceeds.

**Figure 7.**
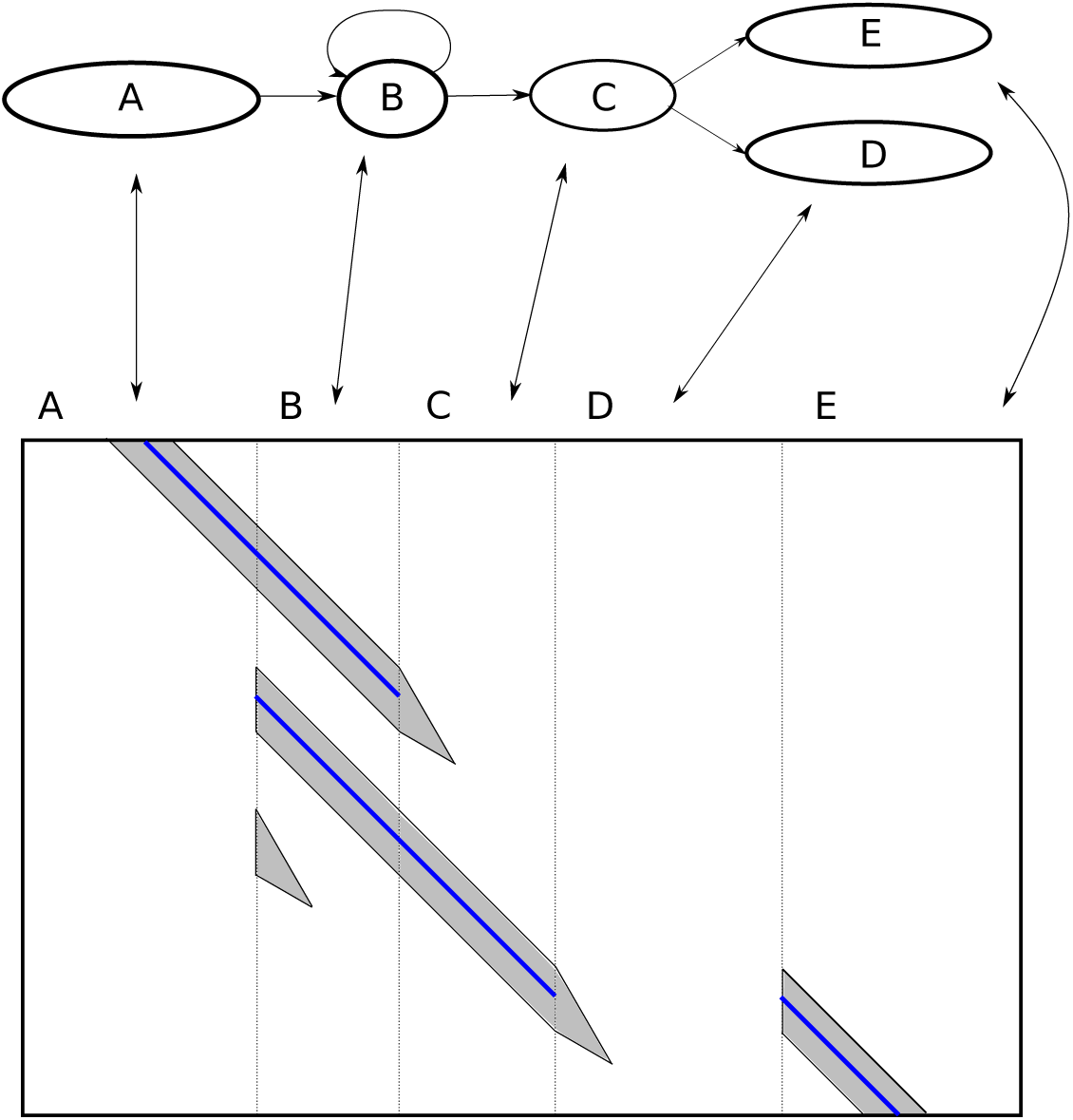
Dynamic score-based banding applied on a graph. Top: an alignment graph. Bottom: The DP matrix for aligning a read to the above graph. The arrows show the correspondence between nodes in the graph and columns in the DP matrix. The dotted lines separate the nodes. The gray area represents parts of the DP matrix which are calculated, and the parts in the white area are not calculated. At each fork, the band spreads to all out-neighbors. The score-based banding implicitly limits the exploration of the alternate paths; as the scores in the alternate paths become worse than the optimal path, the explored part shrinks until finally the exploration stops completely. The blue line shows the backtrace path.

Due to the bitvector-based calculation, the implementation is slightly different from the theoretical description above. The band is defined over blocks in the DP matrix (see Figure 9) instead of individual cells in the DP matrix. In addition, a block’s minimum score is compared to the minimum score in the last row of a 64-row slice.

Since *b* represents a score difference, the score guarantee is now stronger than in the linear case. The optimal alignment is found as long as the optimal alignment’s score at any row is within *b* of the minimum score of that row. This trivially includes the case that the optimal alignment has *b* errors.

However, the size of the band is no longer bounded by *b*. This means that the score-based banding can lead to an impractically large band in certain cases. Figure 8 shows a subgraph of a human whole genome de Bruijn graph. In this case, including all cells with a low score difference will contain a very large part of the subgraph, increasing both runtime and memory use. To handle these cases, we introduce a second banding parameter, the *tangle effort C*. This determines how much effort the DP extension spends in tangled areas of the graph. As we calculate the DP matrix, we keep track of how many cells have been calculated in the current slice. Once this number grows above *C*, we stop calculating the current slice and move to the next slice. This bounds the runtime in tangled regions. However, this is a heuristic method which depends on the assumption that the correct path will be calculated before false positive paths.

**Figure 8.**
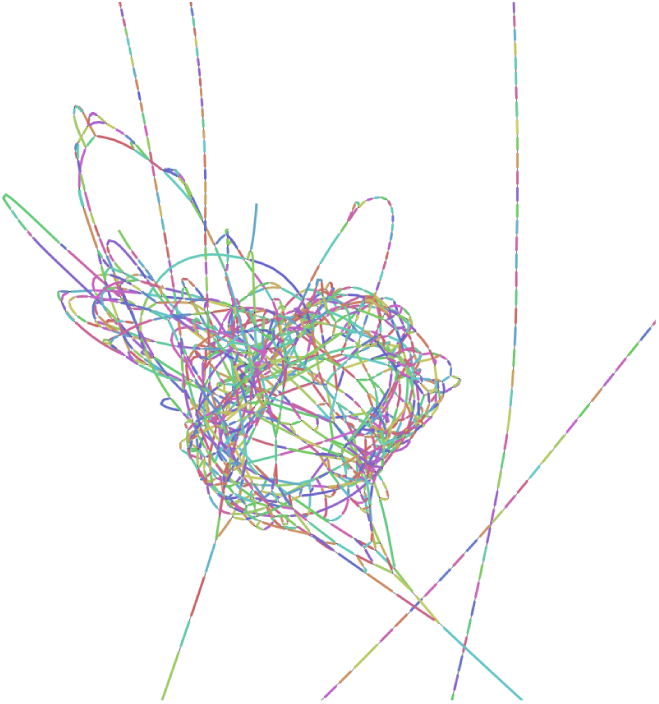
A tangled subgraph of a whole human genome de Bruijn graph. Visualized with Bandage [51].

**Figure 9.**
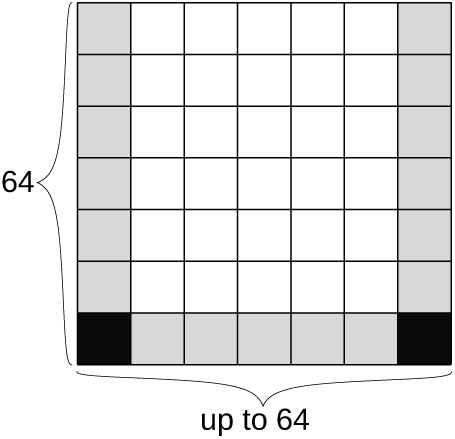
Sparse storage of the DP matrix. Each node is stored in blocks of 64 rows and up to 64 columns. The scores of the corner cells (solid black) are stored explicitly, using 4 bytes per cell. The border cells (gray) are stored with a score difference, using 2 bits per cell. The middle cells (white) are not stored.

In our previous work [23] we used the *minimum changed value* to decide the order in which we calculate the DP matrix. If the parameter *C* is not given, the DP extension uses the minimum changed value as described in the earlier work. However, if the parameter *C* is given, we use a different order, the *minimum changed priority value* of a cell to decide the order. We define the *priority value* of a cell based on the observed error rate of the best alignment so far. With an error rate *e*, a DP cell at row *m* with a score of *k* has a priority value of 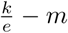 or 64*k - m* if 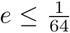. When recalculating a column, the *changed priority value* of a cell is the priority value of a cell in that column which changed, and the *minimum changed priority value* of a column is the minimum of the changed priority values. The intuition is that the priority value of a cell describes “how good” the alignment at a cell is; a value of 0 means as good as the best alignment so far, negative is better than that and positive worse than that. The minimum changed priority value is essentially a greedy heuristic for exploring the most promising paths first. The result is that the minimum changed priority value leads to a higher probability of correctly aligning through a tangle than the minimum changed value when the tangle effort is limited. Without a limit on the tangle effort, using the minimum changed priority value would lead to the scores eventually converging to the same values as the minimum changed value, but the worst case runtime bounds are worse than for the minimum changed value.

### 5.7 Storing a partial DP matrix

In sequence-to-sequence alignment, the banded DP matrix can efficiently be stored as a two-dimensional matrix with 2*b* diagonals, where *b* is the width of the band. However in sequence-to-graph alignment, the banded matrix cannot be stored contiguously due to the non-linear nature of graphs. We conceptually treat the DP matrix as a sparse three-dimensional matrix, with one dimension for node ID, one for node offset and one for read offset.

The implementation stores the DP matrix as a hash table from node IDs to a sparse representation of the alignment between a substring of the read and the sequence of a node. The sparse representation explicitly stores scores at the “bottom corners”, and the score differences between the left, right, and bottom “border cells”. Figure 9 shows an example of this. The middle cells are not stored at all. Instead, the explicitly stored cells allow recalculating the middle cells when needed. This only happens when recalculating cyclic areas, which requires recalculating the middle cells anyway; and during the backtrace, which requires recalculating only the path taken by the backtrace. The sparse representation requires 56 bytes per node, plus memory overhead from the hash table, while using the same data representation that the bit-parallel calculation uses and having no runtime overhead from compression or conversion between different formats. For comparison, the information theoretic lower bound for storing all cells in the DP matrix for one node with optimal compression is 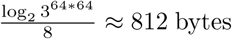 bytes, and storing only the border cells is 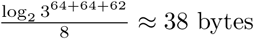 bytes.

### 5.8 Partial alignments

During alignment, we use Viterbi’s algorithm [52] to estimate the correctness at each slice boundary. The observations of the algorithm are the minimum scores at the end of each slice. The two hidden states are *correctly aligned* and *wrongly aligned*. We expect that the correctly aligned state outputs an error rate of 20% and the wrongly aligned an error rate of 50%. These error rates were selected empirically by aligning Oxford Nanopore reads to either the correct or the wrong genomic position. Given the Viterbi estimate, we define a slice as *guaranteed correct* as slices where the backtrace for the wrong state is from the correct state. We also define *guaranteed wrong* slices where the backtrace for the correct state is from the wrong state.

We use the Viterbi estimate to vary the banding parameters. We use two parameters, an initial banding parameter *b* and a ramp banding parameter *B* > *b*. Once the probability of the wrongly aligned state is higher than the probability of the correctly aligned state, we backtrace to the last guaranteed correct slice, switch to the higher ramp banding parameter, and re-align until we have reached the original slice.

We also use the Viterbi estimate to end the alignment. Once we have reached a guaranteed wrong state, the extension can no longer produce anything useful. In this case, we backtrace to the last correct slice and return the partial alignment of the read up to that position.

After extending the seed hits, we are left with a list of partial alignments. We then select a non-overlapping subset of primary and supplementary alignments in a heuristic manner. We greedily pick alignments from longest to shortest, and include an alignment as long as it does not overlap with a previously picked alignment. The primary and supplementary alignments are then written as output. The overlapping alignments are considered secondary and discarded by default, with an optional switch to output secondary alignments as well.

### 5.9 Parallelism

GraphAligner uses a trivial parallelization method by giving each read to a separate thread. Two IO threads handle disk access, and an arbitrary number of worker threads align. One IO thread reads the input reads from a file, and passes them to a single-producer multiple-consumer queue. Each worker thread then takes reads from the queue one at a time and aligns them. Once a worker thread has finished processing a read, it outputs the result into a multiple-producer single-consumer queue. This is then read by the second IO thread which writes the results to a file. The alignment of individual reads is not parallelized.

### 5.10 Experimental setup

All experiments were ran on a computing server with 48 Intel(R) Xeon(R) E7-8857 v2 CPUs and 1.5Tb of RAM. Every program was given 40 threads in the command line invocation. Runtime and memory use was measured with “/usr/bin/time -v” in all experiments.

In the linear comparison experiment, we ran minimap2 with the command “minimap2 -x map-pb -a -t 40”, and BWA with “bwa index” and “bwa mem -x pacbio -t 40”, both corresponding to the recommended parameters for PacBio reads and 40 threads. We ran GraphAligner with “GraphAligner -t 40”, using 40 threads. We used minimap2 version 2.17-r941, BWA version 0.7.17-r1188 and GraphAligner version 1.0.9. The alignment error rates for minimap2 and BWA measured using “samtools stats”. For GraphAligner, the error rate was measured from the output sequence and edit distance fields.

In the variation graph experiment, we “chopped” the graph we gave vg as suggested in the vg documentation, splitting nodes into sub-nodes of at most 32 base pairs. For GraphAligner, we did not chop the graph, and kept the long nodes intact. This is due to the seeding strategy (see Section 5.3), which only finds seeds totally contained inside a node. We ran GraphAligner with parameters “GraphAligner -t 40”, using 40 threads. For vg [16], we first indexed the graph with the command “vg index -t 40 -x graph.xg -g graph.gcsa” using 40 threads, and then aligned with the command “vg map -m long -t 40” as suggested for long read alignment and using 40 threads. We used vg version 1.19.0 and GraphAligner version 1.0.9.

## Competing interests

The authors declare that they have no competing interests.

## Author’s contributions

RM and TM conceived and designed the project. RM implemented GraphAligner. RM ran the experiments. RM and TM wrote the paper.

## Acknowledgements

We thank Rayan Chikhi for advice on how to construct de Bruijn graphs.

## Availability of data and materials

GraphAligner is available from https://anaconda.org/bioconda/graphaligner and https://github.com/maickrau/GraphAligner. Human genome PacBio Sequel data for HG00733 is available from SRA accession SRX4480530 and Illumina from SRA accessions ERR899724, ERR899725, ERR899726. *D. melanogaster* ONT data is available from SRA accession SRR6702603 and Illumina from SRA accession SRR6702604. *E. coli* PacBio data is available from PacBio at https://github.com/PacificBiosciences/DevNet/wiki/E.-coli-Bacterial-Assembly and Illumina data from Illumina at ftp://webdata:webdata@ussd-ftp.illumina.com/Data/SequencingRuns/MG1655/MiSeq_Ecoli_MG1655_110721_PF_R1.fastq.gz and ftp://webdata:webdata@ussd-ftp.illumina.com/Data/SequencingRuns/MG1655/MiSeq_Ecoli_MG1655_110721_PF_R2.fastq.gz

## Funding

MR was funded by the International Max Planck Research School in Computer Science (IMPRS-CS) and acknowledges travel support by the Graduate School for Computer Science Saarbruücken.

## Consent for publication

Not applicable.

## Ethics approval and consent to participate

Not applicable.

## Footnotes

[1] SRA accession SRX4480530

[2] SRA accession SRX4480530

[3] SRA accession SRX4480530

[4] SRA accessions ERR899724, ERR899725, ERR899726

