## Supplementary material for "GraphAligner: Rapid and Versatile Sequence-to-Graph Alignment"

### A. $O(w)$ bitvector merging algorithm

The DP extension algorithm requires merging bitvectors at nodes with an in-degree of at least two. In our previous work Rautiainen *et al.* (2019) we described an $O(\log w)$ algorithm for merging two bitvectors. This was based on finding *difference masks* between the bitvectors, describing indices where one bitvector was greater than the other and vice versa.

We reuse the notation from our previous work. A column $A$ consists of $w$ cells (where $w = 64$ in practice), each of which has an integer value. We notate the value at the $i$'th cell as $S_i^A$. The score difference between two adjacent cells is $S_i^A - S_{i-1}^A \in \{-1, 0, 1\}$. This enables the column to be encoded by two bitvectors $VP^A$ for indices where $S_i^A = S_{i-1}^A + 1$ and $VN^A$ for indices where $S_i^A = S_{i-1}^A - 1$ and a score at end $S_{end}^A = S_{w-1}^A$, which can be used to calculate the score before the first cell $S_b^A = S_{end}^A + popcount(VN^A) - popcount(VP^A)$.

Given two columns $A$ and $B$, we want to find two *difference masks* $M_{A>B}$ and $M_{B>A}$, where $M_{A>B}$ is set in indices where $S^A > S^B$ and $M_{B>A}$ is set in indices where $S^B > S^A$. The idea is to first create a *difference bitvector* of the difference between the two bitvectors, which describes how the score difference $S^A - S^B$ changes at each index. The difference bitvector consists of four bitvectors *twosmaller*, *onesmaller*, *onebigger* and *twobigger*, which correspond to the cases where $(S_i^A - S_i^B) - (S_{i-1}^A - S_{i-1}^B) \in \{-2, -1, 1, 2\}$ respectively. Given the difference bitvector, we find the first index where $S^A - S^B$ increases, and the first index where the $S^A - B^A$ decreases. Assume without loss of generality that we have two bitvectors with the same score before the first cell $S_b^A = S_b^B$, the first (least significant) index $i$ where the difference $S^A - S^B$ increases and the first index $j$ where the difference $S^A - S^B$ decreases, with $i < j$. Then, *regardless of the scores elsewhere in the bitvector*, $S_x^A > S_x^B$ in indices $i \leq x < j$. Using this idea, we can create an algorithm which calculates at least one index of the difference masks in each iteration. The algorithm repeatedly takes the smallest indices $i$ and $j$ where the difference increases and decreases, sets the difference masks in $[i, j)$, then unsets the difference bitvectors at the indices $i$ and $j$. Once the smaller bitvector has no more set bits, the remaining masks are set according to whether the bigger bitvector has bits left, and similarly vice versa. This results in an algorithm with a runtime of $O(w)$.

The above assumed that the scores before the first cells are equal, that is, $S_b^A = S_b^B$. This is not always true. To solve this corner case, we run a preprocessing step where we pretend that the difference bitvectors have extra bits set "before" the first cell, and run the bit unsetting described above on those extra bits. The source code of the merge operation in C++ is below.

```
//Word is an unsigned integer of arbitrary width, in practice uint64_t
template <typename Word>
std::pair<Word, Word>
differenceMasks(Word VP_A, Word VN_A, int Sb_A, Word VP_B, Word VN_B, int Sb_B)
{
Word ASmaller = 0;
Word BSmaller = 0;
//unset bits which are common in both bitvectors
//that is, both bitvectors increase or decrease
Word VPcommon = ~(VP_A & VP_B);
Word VNcommon = ~(VN_A & VN_B);
VP_A &= VPcommon;
VN_A &= VNcommon;
VP_B &= VPcommon;
VN_B &= VNcommon;
//difference bitvectors describing the score change at each index
Word twosmaller = VN_A & VP_B; //A is two smaller
Word onesmaller = (VP_B & ~VN_A) | (VN_A & ~VP_B);
Word onebigger = (VP_A & ~VN_B) | (VN_B & ~VP_A);
Word twobigger = VN_B & VP_A; //A is two bigger
onebigger |= twobigger;
onesmaller |= twosmaller;
//preprocessing: handle the cases where Sb_B != Sb_A
if (Sb_B > Sb_A)
{
 for (int i = 1; i < Sb_B - Sb_A; i++)
 {
  Word leastSignificant = onebigger & ~(onebigger - 1);
  onebigger ^= (~twobigger & leastSignificant);
  twobigger &= ~leastSignificant;
  if (onebigger == 0)
  {
   return std::make_pair(-1, 0);
  }
 }
 Word leastSignificant = onebigger & ~(onebigger - 1);
```

```
 ASmaller |= leastSignificant - 1;
 onebigger ^= (~twobigger & leastSignificant);
 twobigger &= ~leastSignificant;
}
else if (Sb_A > Sb_B)
{
 for (int i = 1; i < (Sb_A - Sb_B); i++)
 {
  Word leastSignificant = onesmaller & ~(onesmaller - 1);
  onesmaller ^= (~twosmaller & leastSignificant);
  twosmaller &= ~leastSignificant;
  if (onesmaller == 0)
  {
   return std::make_pair(0, -1);
  }
 }
 Word leastSignificant = onesmaller & ~(onesmaller - 1);
 BSmaller |= leastSignificant - 1;
 onesmaller ^= (~twosmaller & leastSignificant);
 twosmaller &= ~leastSignificant;
}
//unset bits one at a time
for (int i = 0; i < sizeof(Word)*8; i++)
{
 if (onesmaller == 0)
 {
  if (onebigger == 0) break;
  Word leastSignificant = onebigger & ~(onebigger - 1);
  BSmaller |= -leastSignificant;
  break;
 }
 if (onebigger == 0)
 {
  assert(onesmaller != 0);
  Word leastSignificant = onesmaller & ~(onesmaller - 1);
  ASmaller |= -leastSignificant;
  break;
 }
 Word leastSignificantBigger = onebigger & ~(onebigger - 1);
 Word leastSignificantSmaller = onesmaller & ~(onesmaller - 1);
 assert((onebigger & leastSignificantBigger) != 0);
 assert((onesmaller & leastSignificantSmaller) != 0);
 assert(leastSignificantSmaller != leastSignificantBigger);
 assert(leastSignificantSmaller != 0);
 assert(leastSignificantBigger != 0);
```

```
  if (leastSignificantBigger > leastSignificantSmaller)
  {
   ASmaller |= leastSignificantBigger - leastSignificantSmaller;
  }
  else
  {
   BSmaller |= leastSignificantSmaller - leastSignificantBigger;
  }
  onebigger ^= (~twobigger & leastSignificantBigger);
  twobigger &= ~leastSignificantBigger;
  onesmaller ^= (~twosmaller & leastSignificantSmaller);
  twosmaller &= ~leastSignificantSmaller;
 }
 assert((ASmaller & BSmaller) == 0);
 assert(onesmaller == 0 || onebigger == 0);
 return std::make_pair(ASmaller, BSmaller);
 }
```
